# Extensive Soma-Soma Plate-Like Contact Sites (Ephapses) Connect Suprachiasmatic Nucleus Neurons

**DOI:** 10.1101/2022.09.09.507192

**Authors:** Mark É. Czeisler, Yongli Shan, Richard Schalek, Daniel R. Berger, Adi Suissa-Peleg, Joseph S. Takahashi, Jeff W. Lichtman

**Affiliations:** Department of Molecular and Cellular Biology, Harvard University, Cambridge, MA, 02138, USA; Department of Neuroscience, Peter O’Donnell Jr. Brain Institute, University of Texas Southwestern Medical Center, Dallas, TX 75390-9111, USA; Howard Hughes Medical Institute, University of Texas Southwestern Medical Center, Dallas, TX 75390-9111, USA

**Keywords:** circadian rhythms, suprachiasmatic nucleus (SCN), serial electron microscopy, ultrastructure, mouse, ephaptic coupling

## Abstract

The hypothalamic suprachiasmatic nucleus (SCN) is the central pacemaker for mammalian circadian rhythms. As such, this ensemble of cell-autonomous neuronal oscillators with divergent periods must maintain coordinated oscillations. To investigate ultrastructural features enabling such synchronization, 805 coronal ultrathin sections of mouse SCN tissue were imaged with electron microscopy and aligned into a volumetric stack, from which selected neurons within the SCN core were reconstructed *in silica*. We found that clustered SCN core neurons were physically connected to each other via multiple large soma-to-soma plate-like contacts. In some cases, a sliver of a glial process was interleaved. These contacts were large, covering on average ∼21% of apposing neuronal somata. It is possible that contacts may be the electrophysiological substrate for synchronization between SCN neurons. Such plate-like contacts may explain why synchronization of SCN neurons is maintained even when chemical synaptic transmission or electrical synaptic transmission via gap junctions is blocked. Such ephaptic contact-mediated synchronization among nearby neurons may therefore underlie the wave-like oscillations of circadian core clock genes and calcium signals observed in the SCN.

**Significance:** Three-dimensional reconstruction of SCN tissue via serial electron microscopy revealed a novel structural feature of SCN neurons that may account for inter-neuronal synchronization that persists even when usual mechanisms of neuronal communication are blocked. We found that SCN core neurons are connected by multiple soma-soma contact specializations, ultrastructural elements that could enable synchronization of tightly packed neurons organized in clustered networks. This extensive network of plate-like soma-soma contacts among clustered SCN neurons may provide insight into how ∼20,000 autonomous neuronal oscillators with a broad range of intrinsic periods remain synchronized in the absence of ordinary communication modalities, thereby conferring the resilience required for the SCN to function as the mammalian circadian pacemaker.

## Introduction

In mammals, the suprachiasmatic nucleus (SCN) of the hypothalamus is the pacesetting circadian clock that drives circadian oscillations governing physiology and behavior, including those driving daily cycles of body temperature, endocrine, hepatic, cardiac, pulmonary, immune and digestive functions, sleeping, eating, and activity in mammals (Hastings et al., 2007). The SCN is synchronized to the 24-hour day by the light-dark cycle through a monosynaptic pathway that links intrinsically photosensitive ganglion cells (ipRGCs) directly to SCN neurons in the ventrolateral region of the SCN (the SCN core), which contains neurons with the capacity for light-induced gene expression (Schmidt et al., 2011). The dorsolateral region of the SCN (the SCN shell) integrates photic information and other time cues conveyed by SCN core neurons to generate circadian output rhythms, which are then projected to the rest of the brain and body (Shan et al., 2020).

Circadian activity rhythms are driven by a transcriptional-translational feedback loop of circadian clock genes. The precision of the clock is dramatically improved by synchronization of individual intact SCN pacemaker neurons, can persist at a stable near-24-hour period, even in the absence of light-based cues. Remarkably, Schwartz *et al*. (1987) showed that circadian oscillations in the SCN persisted with a stable near-24-hour period for weeks, even when tetrodotoxin (TTX) was continuously infused into the SCN, blocking sodium-dependent chemical synapse transmission of input pathways for entrainment, intracellular communications for synchronization, and output pathways for expression of circadian rhythms (Schwartz et al., 1987). Thus, it was surprising when Welsh *et al*. (1995) discovered that individual, dissociated SCN neurons exhibit highly variable autonomous circadian periods, ranging from 22 to 30 hours (Welsh et al., 1995; Liu et al., 2007; Ko et al., 2010). Generating coherent circadian rhythms at the tissue-wide level of the SCN from noisy component cells requires coordination of the independent cell-autonomous oscillators.

Despite an increasing understanding of the role of the SCN core—specifically neuropeptide transmitters (vasoactive intestinal polypeptide, calretinin, neurotensin, and gastrin releasing peptide) and GABA—in the synchronization of circadian output rhythms, mechanisms governing core-localized inter-neuronal communication have yet to be elucidated. Indeed, in contrast to the coordinated oscillations of the bilateral SCN under 12-hour:12-hour light:dark conditions, the left and right SCN oscillate in antiphase under constant light conditions (de la Iglesia et al., 2000). This achievement of a “split” brain in the absence of a surgical bisection suggests that light input strongly drives entrainment of the SCN hemispheres that would otherwise inhibit each other, further supporting the necessity of local synchronization to maintain these rhythms.

It has since been shown that while TTX reduces the amplitude of these oscillations within the SCN (Yamaguchi et al., 2003; Buhr et al., 2010), they remain coherently synchronized regionally within an *ex vivo* SCN slice (Enoki et al., 2012), suggesting that both chemical synaptic coupling and non-chemical synaptic coupling between SCN neurons are involved in maintaining synchronization within regional clusters of SCN core neurons. Enoki et al. (2012) applied carbenoxolone (CBX), a gap junction blocker, and found that synchronization of SCN neurons persisted. In 2017, Diemer *et al*. evaluated gap junctions using a different approach, as they generated a Connexin-36 double-knockout mouse, which demonstrated similarly synchronized neurons (Diemer et al., 2017).

Thus, synchronization of SCN neurons with divergent periods persists even when chemical or gap junction mediated electrical synapses are blocked. These findings leave open the possibility that synchronization is based on some non-synaptic mechanism. Therefore, to investigate if there might be an alternative structural basis for SCN synchronicity, we generated a 3-dimensional reconstruction from serial electron micrographs of SCN neurons (an SCN connectome) to characterize the structural and functional architecture of a cluster of neurons in the SCN core at nanometer resolution.

## Materials and Methods

### Experimental Procedures

All animal studies and Materials and methods were in accordance with UT Southwestern Medical Center guidelines for animal care and use. The mouse used was housed at UT Southwestern Medical Center on a 12:12 LD cycle. A ∼300 × 500 × 1,000-μm volume encompassing the SCN, third ventricle, and part of the optic chiasm was isolated from a 4-month-old C57BL/6N *wildtype* male mouse that was euthanized at 16:00 [Zeitgeber time (ZT) 10] via a euthasol injection and perfused transcardially with a fixative solution containing 2% paraformaldehyde and 2.5% glutaraldehyde in a CaCl_2_ (3mM), sodium cacodylate (0.15M) buffer. The tissue was shipped on ice to Harvard University for washing and staining via the reduced osmium-thiocarbohydrazide-osmium protocol (exposure to 4% osmium tetroxide-potassium ferrocyanide for 60 minutes, 1% thiocarbohydrazide for 30 minutes, then 2% osmium tetroxide for 60 minutes with extended washes using double-distilled H_2_O (ddH_2_O) in between) (Tapia et al., 2012). The tissue was incubated overnight at 4°C in 1% uranyl acetate and then dehydrated using alcohols (5-minute washes in 50%, 70%, 90%, 100%, and 100% ethanol in ddH_2_O, then 2 × 5-minute washes in 100% propylene oxides). Finally, propylene oxide was replaced step-wise with 812 Epon resin (60 minutes in 3:1 propylene oxide/resin, overnight in 1:1 propylene oxide/resin, and incubated for 72 hours at 60°C in 0:1 propylene oxide/resin), curing the resin into a plastic block encapsulating the sample. 805 consecutive 50-nm ultrathin serial sections were cut and collected onto a 50-μm thick carbon-coated Kapton tape reel from the embedded tissue block using an automated tape-collecting ultra-microtome (ATUM) (Hayworth et al., 2014) (Fig. 1A).

**Figure 1.**
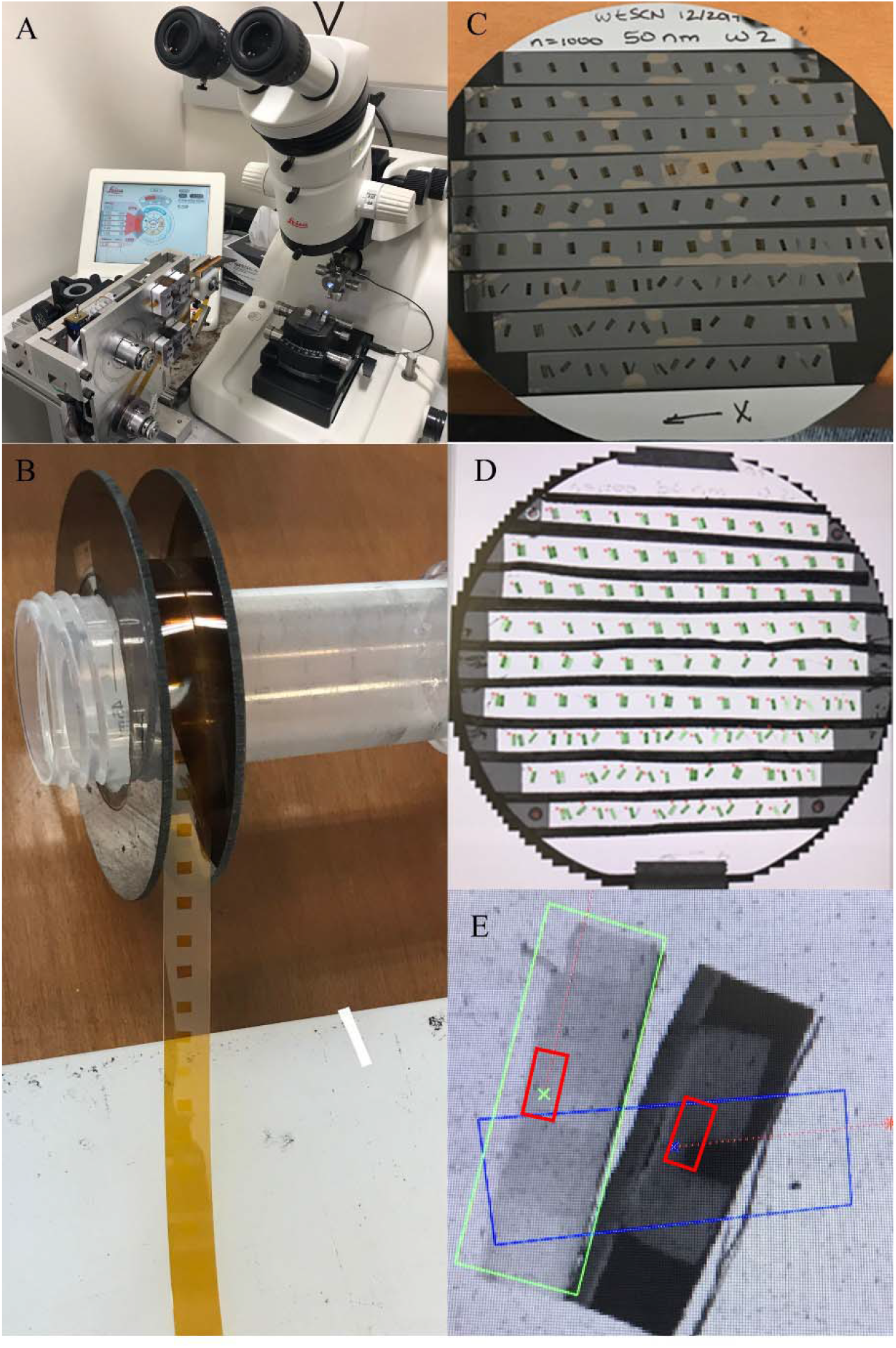
Serial Sectioning, Identifying and Labeling a Region of Interest (ROI) for High-Resolution Electron Microscopical Imaging. (A) The automated tape-collecting ultramicrotome (ATUM). The bottom reel of the ATUM contains tape that is fed into the knife boat of a diamond knife mounted on a commercial ultramicrotome. A diamond knife boat is filled with water and a diamond knife cuts serial ultrathin sections from tissue embedded in a plastic block. Sections float on the surface of the water in the knife boat until they adhere to the moving tape. (B) Kapton tape roll containing sections collected with the ATUM. (C) Section-containing tape was transferred from the roll to silicon wafers. (D) An optical image of one of five tape-containing silicon wafers contains ∼200 sections from the mouse SCN. Optical images were used to automate the EM imaging with reference locations for each section. The red rectangle encloses two consecutive sections shown in (E). (E) Optical image of two sections containing mouse SCN tissue. The green box shows completed manual section spatial-labeling for semi-automatic EM imaging. Red boxes in Panel 3D indicate the 330×150-μm SCN region of interest that was manually labeled for high-resolution imaging.

Each ∼300 × 500 × 0.05-μm section contained the entire region of interest (ROI) coronally and the stack of sections cumulatively represented 40 μm of the ∼300-μm (13.3%) depth of the SCN (Fig. 1B). The section-containing tape was manually cut into 5-9 cm strips and transferred to 10-centimeter diameter silicon wafers. Sections were stained with 4% uranyl acetate for 3 minutes, washed, and then stained with 4% lead citrate for 3 minutes (Fig. 1C). A 330 × 180-μm coronal region of interest (Fig. 1D) was selected on each section for imaging at high resolution with the scanning electron microscope.

Images of the ROI were acquired from each section through automated imaging commands on ATLAS software (Fibics) connected to a Sigma scanning electron microscope (Carl Zeiss). The electron beam was set to 1.7 kilovolts to use secondary electron detection at a working distance of 3.4 mm and a dwell time of 200 nanoseconds/pixel. Images were acquired at 4 nanometer/pixel resolution, with each image tile containing 16,384 pixels x 16,384 pixels. A mosaic of 18 tiles (6 transverse x 3 longitudinal tiles) was used to capture the ROI with a 6% tile overlap to facilitate stitching between tiles. Tiles within an individual section were stitched to create a single image of each section, then consecutive sections were aligned into a 3-dimensional stack using the elastic alignment method (Saalfeld et al., 2012).

Eight SCN neurons were identified and segmented (labeled) using a computer-assisted manual volume annotation and segmentation tool (VAST) (Berger et al., 2018). Each neuron was arbitrarily assigned a unique color for labeling, which was painted as an overlaid image layer (segmentation layer) superimposed on the EM data (Fig. 2A-B, Mult. 1 [<u>view online</u>]). To characterize appositions between SCN core neurons, a second set of reconstructions restricted segmentation to only the membrane of each neuronal soma. Areas of soma membrane not in direct contact with other neuronal somata were labeled the same color as the original reconstruction, while regions of direct soma-soma plate-like contact sites were assigned a new, previously unused color. Volumetric objects generated from compiling annotations throughout the segmentation layer containing labeled neurons were exported from VAST to 3ds Max (Autodesk, Inc.) to generate comprehensive 3D renderings, adjust smoothing and lighting, and estimate surface area and volume statistics (Fig. 2C).

**Figure 2.**
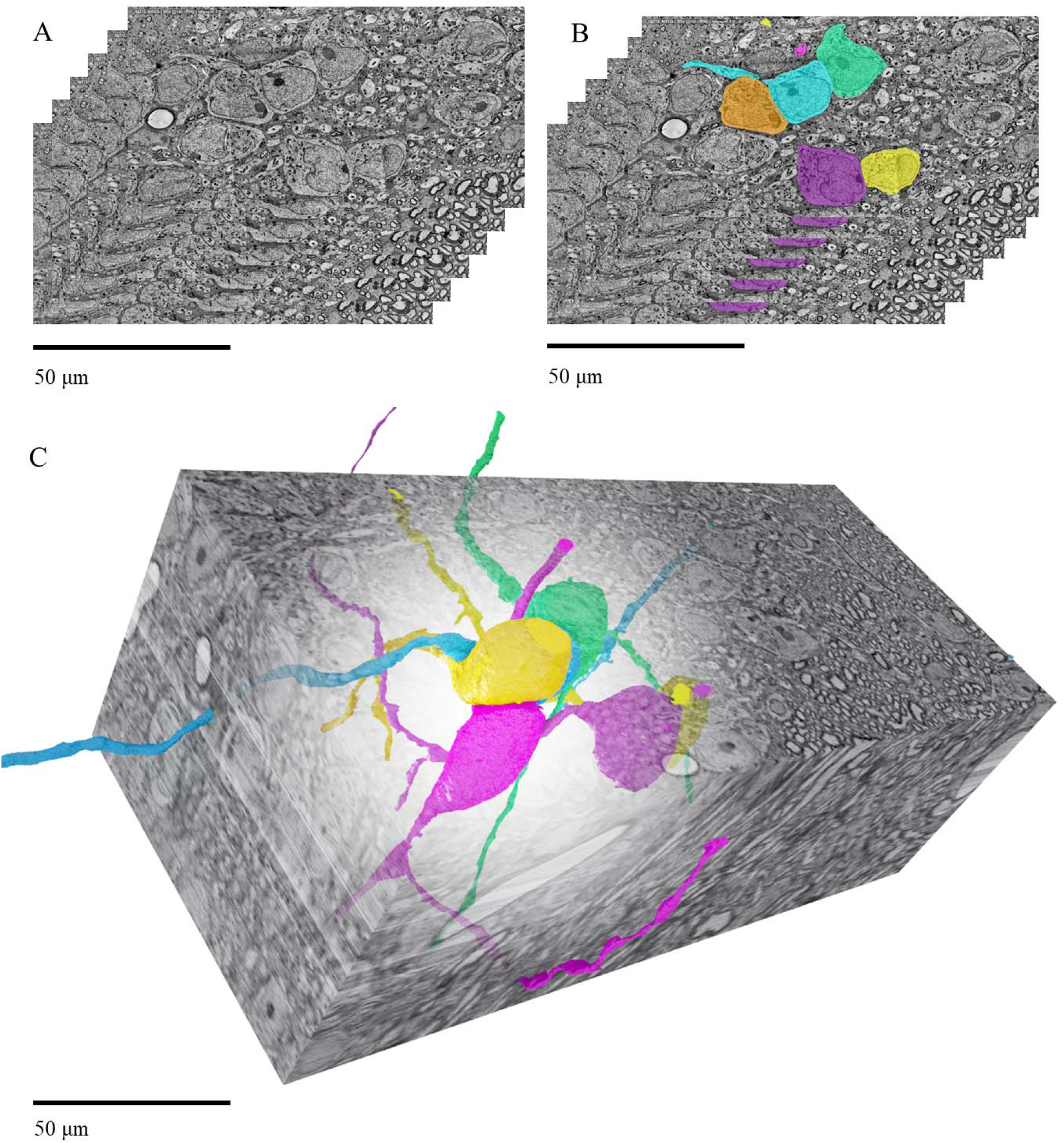
Reconstructing Neurons from Electron Microscopically Imaged Serial Sections. (A) 6 of the 805 consecutive 50-nm serial EM sections that were acquired, aligned, and loaded into Volume Annotation and Segmentation Tool (VAST) for manual segmentation. (B) The same serial EM sections shown in (A) were color-labeled to distinguish individual neurons and processes through a vertical stack of aligned images. (C) Color-labeling performed in VAST on each section in the volume were rendered into a digitized 3-dimensional reconstruction of the neurons using 3ds Max. Adobe Photoshop was used to achieve the transparent effect in (C).

VastTools was used to calculate the total surface area and volume measurements for each of the reconstructed segments. Because VastTools sums the area of generated mesh triangles for the object, it is possible that the calculated area was greater than that of the true object due to the surface roughness. Therefore, surface area and volume statistics were also obtained in 3ds Max after an auto-smoothing modifier was applied to smooth edges from tracing by meshing adjacent faces if the angle between them were less than the threshold angle (30°). The reported surface area and volume statistics are an average of the two methods. In-depth explanations of these features are described in VAST user manual, which is also available for download from the VAST webpage.

Finally, for an expanded, exploratory view of the SCN ultrastructure including regions outside of the ROI, vertical stacks of subsets of the 805 sections from randomly selected areas across both hemispheres and both the SCN shell and core were compiled into videos absent segmentation.

## Results

A digital dataset containing 805 50-nm serial sections representing a 272 × 120 × 40.25-μm volume (1,313,760 μm^3^) of SCN tissue was acquired, stitched and aligned. The resulting data had a 4-nm-per-pixel in plane resolution (see methods for detail; Fig. 3). Eight neurons were reconstructed from this dataset: six adjacent neurons with somata in the SCN core, and two neurons with somata in the optic chiasm and processes extending into the SCN core. A three-dimensional render of the six neurons (color-labeled in blue, yellow, green, orange, pink, or purple) positioned in the SCN core (Fig. 4) revealed extensive loci of soma-soma plate-like contact sites between these neighboring SCN neurons (Mult. 2 [<u>view online</u>]). In some cases, slivers of glial cells that were largely devoid of cytoplasmic material occupied the interstitial space between two closely apposed somata within the soma-soma contact areas. In a few cases, neuronal somata were in direct contact with one another. The estimated mean surface area of the six clustered SCN neuronal somata was 853 μm^2^ ± 146 μm^2^ (SD), and each soma had 1-6 large soma-soma plate-like contact sites that together occupied an average of ∼21% of their somatic surface (Fig. 5, Table). Of the 23 plate-like contact sites on the six neurons constructed within the neuronal cluster, one lacked glial membrane intrusion, whereas 22 had glial membrane intercalations but these were devoid of cytoplasm and occupied only some part of the soma-soma plate-like contact sites.

**Figure 3.**
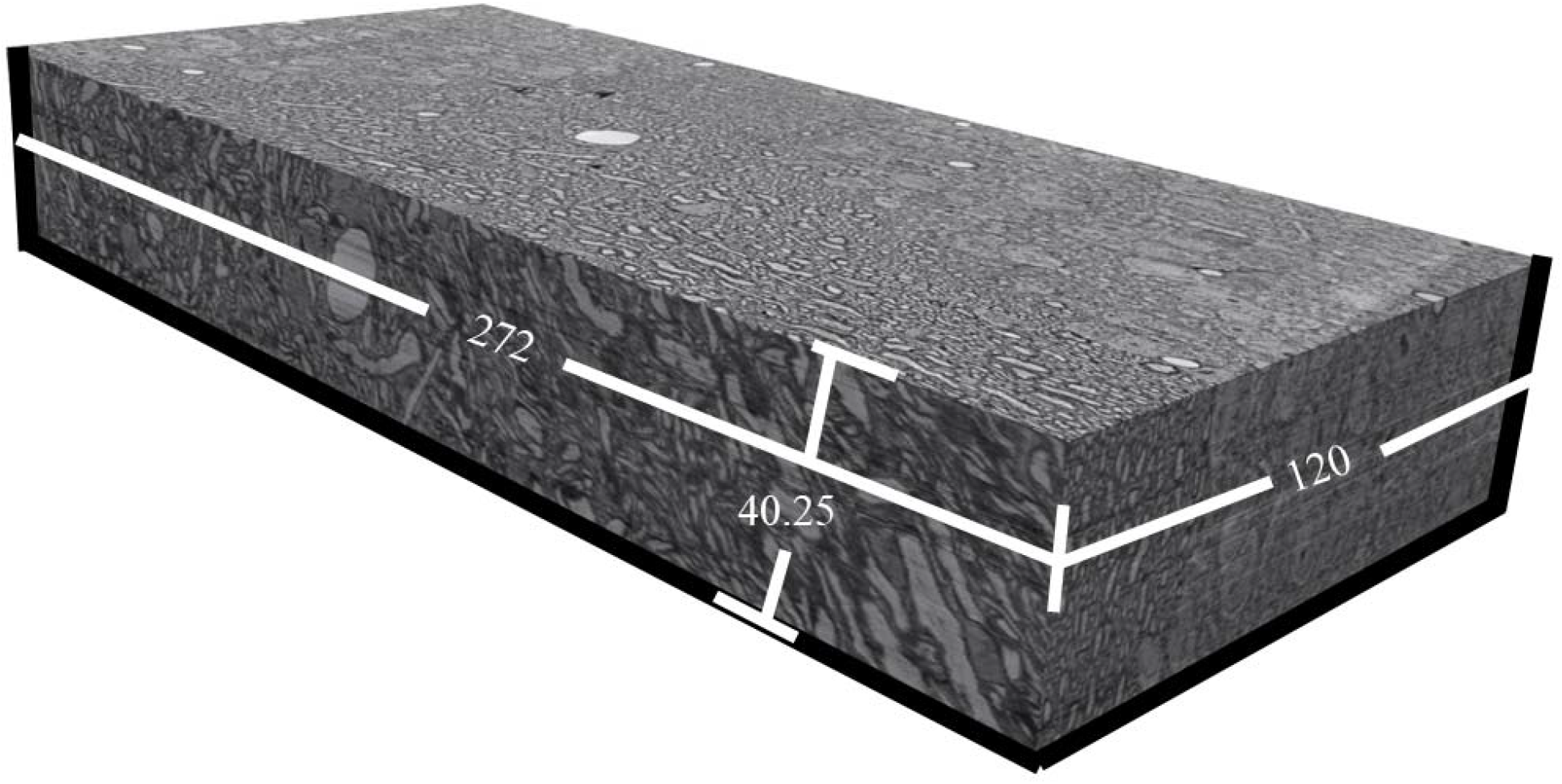
High-Resolution SCN EM Image Block. The entire digitized block of 805 stitched, aligned, and traceable ultrathin serial sections amounted to a volume of 272 × 120 × 40.25-μm (1,313,760-μm^3^).

**Figure 4.**
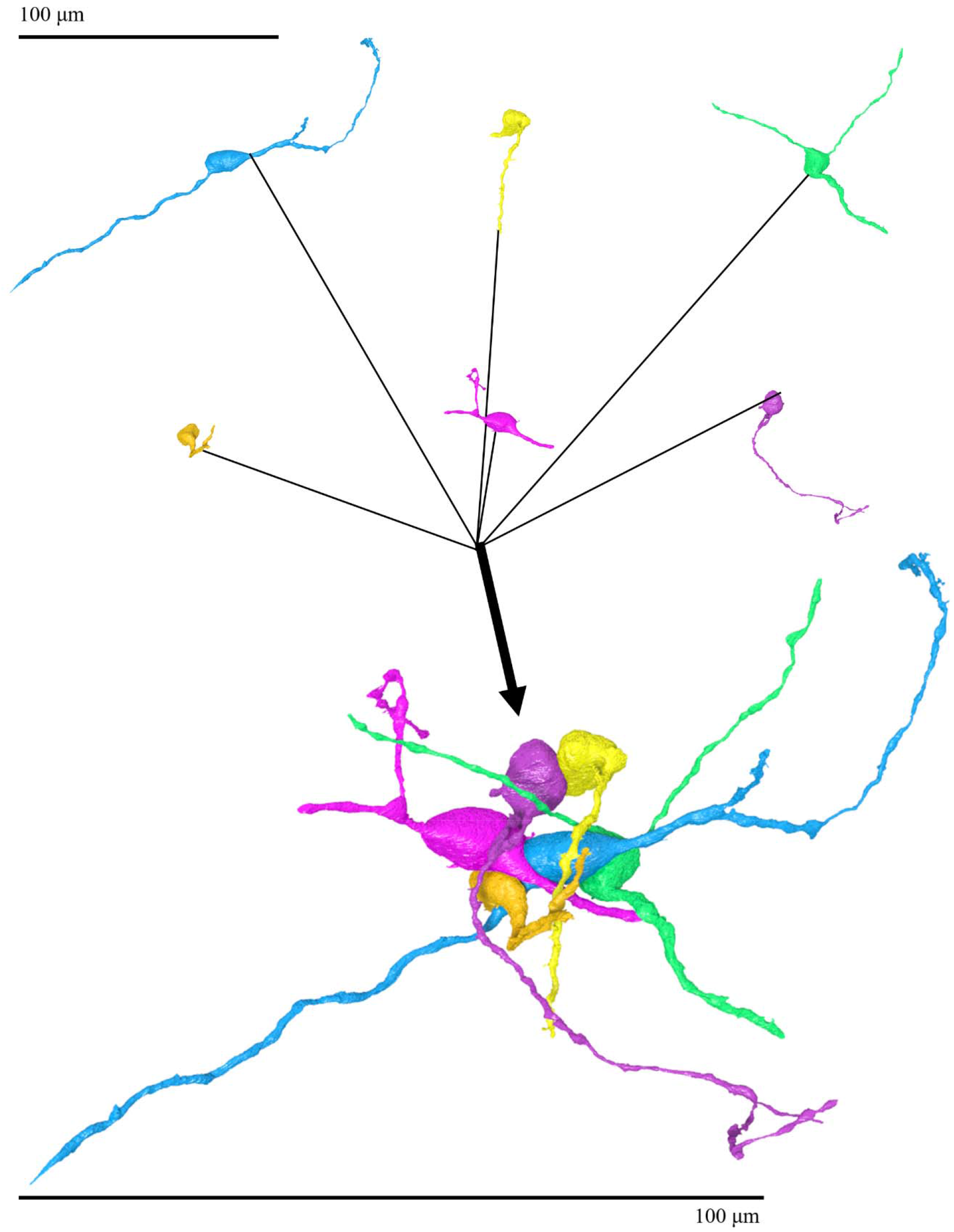
Color-labeled Clustered SCN Core Neurons. Above, each of the six reconstructed SCN core neurons is shown in isolation. Below, all six neurons are shown as they were positioned in the SCN. Note the extensive contact plates between adjacent neurons. Each neuron retains its relative size and orientation between the isolated and merged perspectives. The scale bars represent 100 µm.

**Figure 5.**
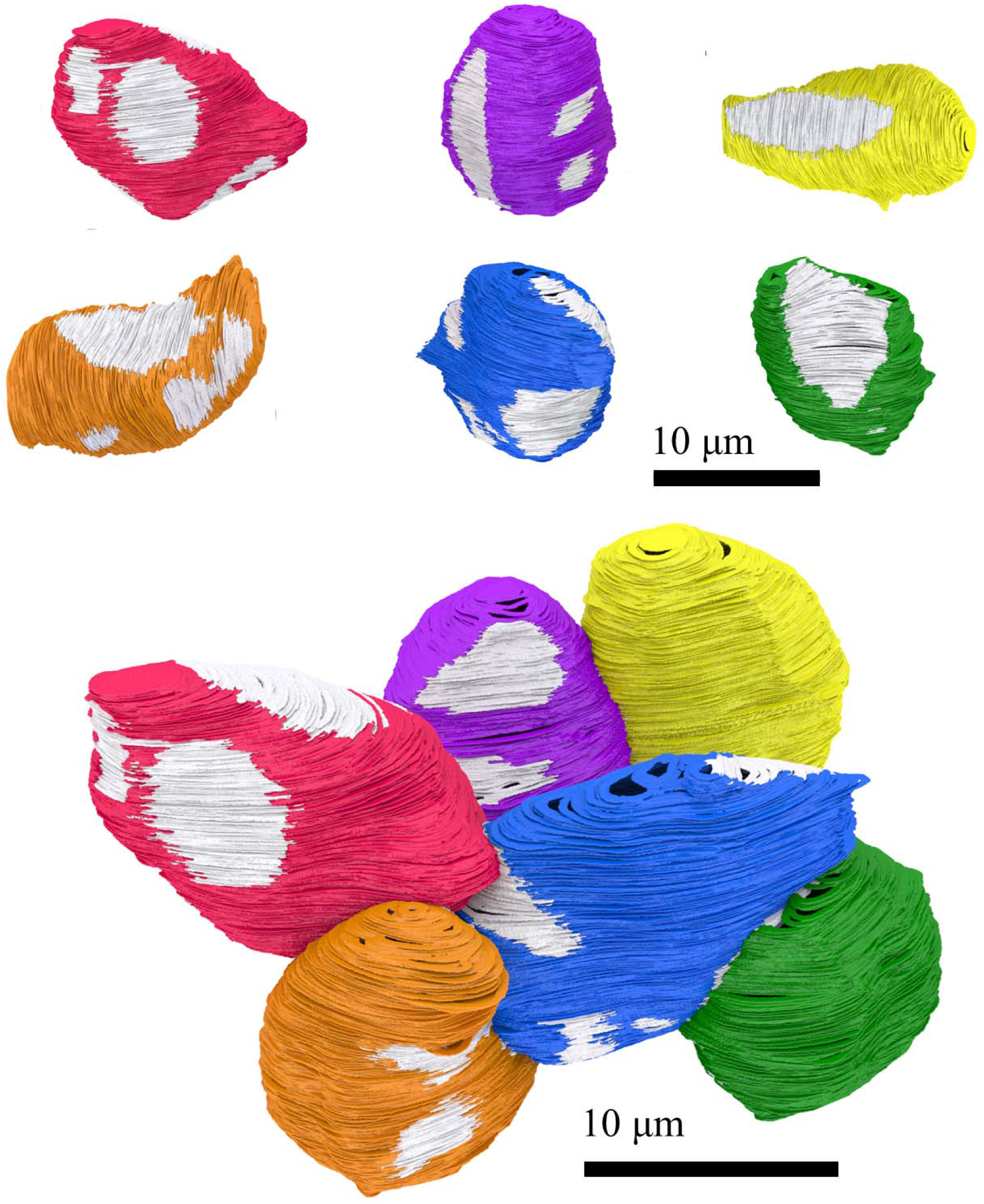
Ephaptic Contact Plates Between SCN Core Neurons Color-labeled. Each of the six neurons retained its original base color-label from the initial reconstruction (blue, yellow, green, orange, pink, and purple). Surfaces of ephaptic contact plates between each neuron and an adjacent soma was labeled in a different, previously unused color (magenta, dark green, tan, lime green, pale green, and dark red). (A-F) Each neuron is shown in isolation, rotated to reveal its most extensive ephapse. (G) All six core neurons are shown as positioned in the SCN core.

**Table.** SCN Core Neuronal Somatic Surface Area.

| Reconstructed Neuronal Somata | Somatic Surface Area ($\mu\text{m}^2$) | Total Ephaptic Plate Surface Area ($\mu\text{m}^2$) | Somatic Surface Area Occupied by an Ephaptic Plate (%) |
| --- | --- | --- | --- |
| Core Blue | 1038.3 | 284.8 | 27.4 |
| Core Yellow | 695.3 | 58.9 | 8.5 |
| Core Green | 726.4 | 103.1 | 14.2 |
| Core Orange | 963.0 | 186.8 | 19.4 |
| Core Pink | 990.8 | 313.7 | 31.7 |
| Core Purple | 704.5 | 125.4 | 17.8 |
| Core Neuronal Somata Mean | 853.1 | 178.8 | 21.0 |

The estimations reported are an average of the surface area calculations of the segments color-labeled with unique colors to distinguish between membrane surface areas that are in direct somatic-ephaptic contact with an adjacent neuron and areas that are not in direct somatic-ephaptic contact. Note that this does not include other ephaptic contact (soma-dendritic, soma-axonal, dendro-dendritic, axo-axonal) which was also visibly present. An identifier for each neuron (Column 1), ephaptic contact surface area (Column 2), total somatic surface area (Column 3) and ephaptic contact surface area as a percentage of the total somatic surface area (Column 4) are reported for each soma in the SCN core neuronal cluster.

The EM images revealed what may be intercellular membrane specializations along portions of some soma-soma plate-like contact sites especially at sites between neurons without interleaved glial cell membranes. Small clusters of electron-dense material were observed along the extracellular face of the contacting membranes, the intensity of which sometimes obscured the narrow soma-soma cleft (Fig. 6). In some cases, there were consecutive, equally spaced punctate densities resembling a zipper along adjacent perikaryal areas (Fig. 9).

**Figure 6.**
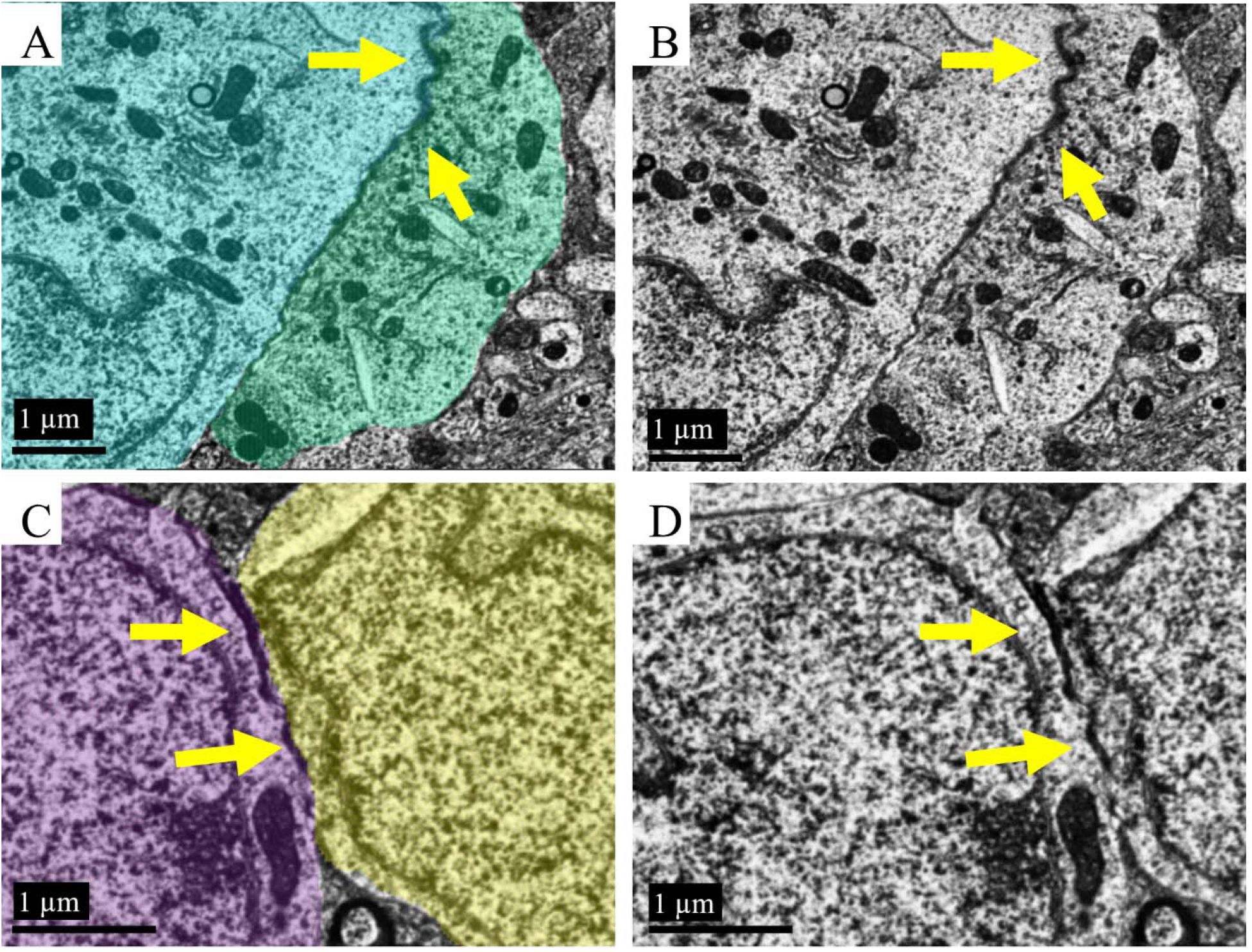
Electron-Dense Deposits Along Ephaptic Contact Plates. Yellow arrows point to electron-dense regions, which are localized to a subset of ephaptic contact plates. (A, C) Single-section examples of dark patches between pairs of segmented and reconstructed cells; such electron-dense, dark patches were common in the volume. (A) shows an ephapse between the SCN core neurons assigned blue and green. (C) shows an ephapse between the SCN core neurons assigned purple and yellow. (B, D) The same frame as shown in (A, C) respectively, without the segmentation color layer offers a clearer view of the dense regions. These electron-dense areas within the ephapses are consistent with those observed using EM when there are gap junctions between cells. Scale bars represent 1 µm.

In addition to 21% of the somatic surface being occupied by soma-soma contact, in some cases, there were soma-dendrite plate-like contact sites between neurons. Three of the six neurons reconstructed with cell bodies in the SCN core had dendrites extending into the optic chiasm, which received synapses deep in the optic chiasm. There were also neuronal cell bodies interspersed among myelinated axons within the optic chiasm. Despite their proximity, the two reconstructed cells with somata embedded in the optic chiasm exhibited no perikaryal contacts.

Finally, examination of six columnar vertical serial EM image stacks from within the SCN block reveal that most neuronal somas that were identified within a frame were closely apposed to at least one other neuronal soma (Fig. 7, Mult. 4-10). These clusters of neuronal somata are separated from other clusters by areas of neuropil.

**Figure 7.**
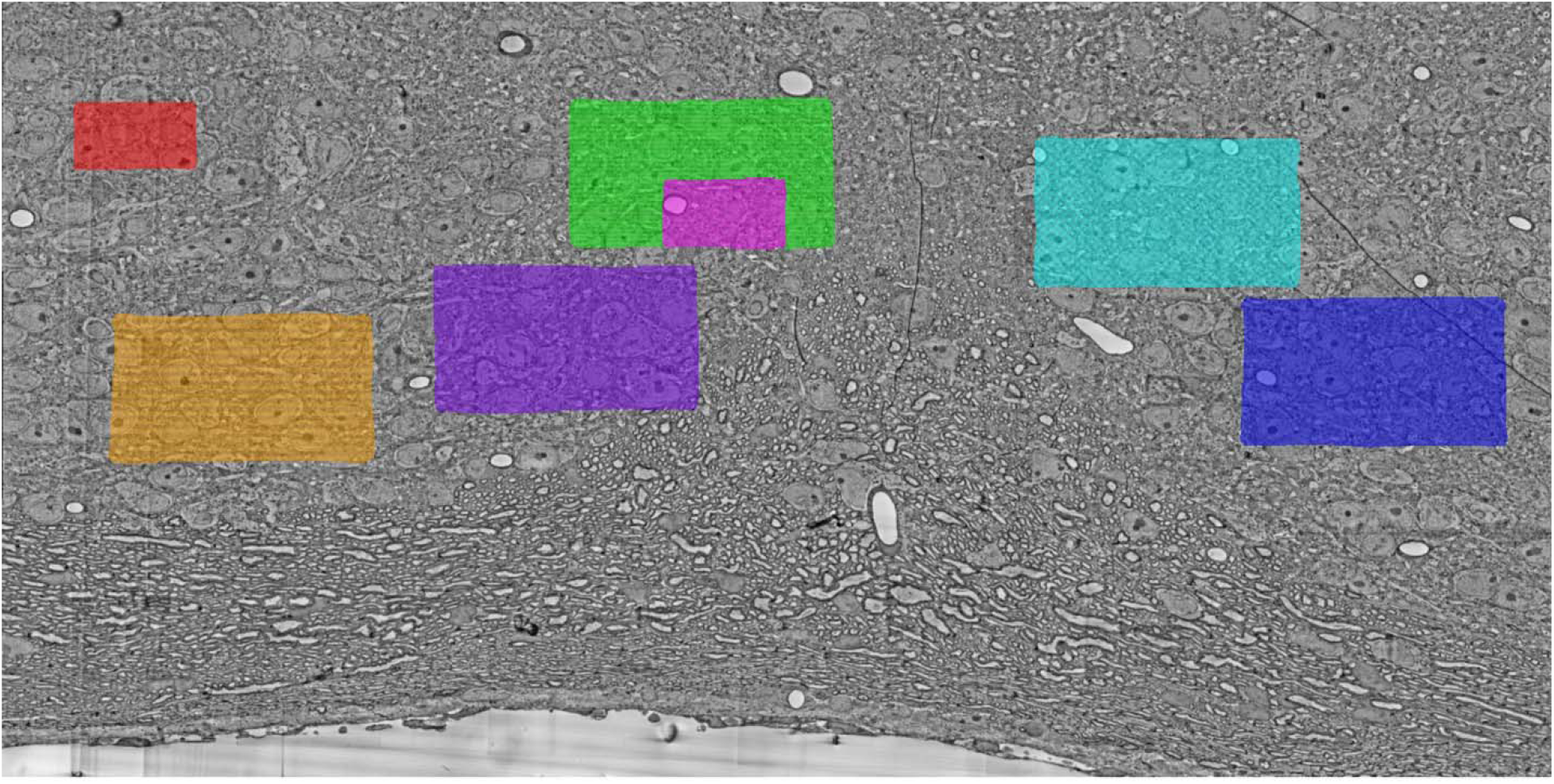
Location of Randomly Selected Regions for Columnar Spacelapse Videos. Video 4 corresponds to the orange window-of-view. Video 5 corresponds to the green window-of-view. Video 6 corresponds to the turquoise window-of-view. Video 7 corresponds to the purple window-of-view. Video 8 corresponds to the blue window-of-view. Video 9 corresponds to the red window-of-view. Video 10 corresponds to the magenta window-of-view.

## Discussion

Three-dimensional reconstructions of core suprachiasmatic nucleus neurons reveal extensive soma-soma plate-like contact sites between adjacent neurons (Mult. 3 [<u>view online</u>]). This neuroanatomic feature, identified by compiling consecutive EM images of ultrathin SCN sections, could account for the persistence of mutual synchronization of the cell-autonomous neuronal oscillators comprising the SCN core that persists when either chemical or electrical synaptic coupling is blocked (Schwartz et al., 1987; Edgar et al., 2012; Diemer et al., 2017). It had previously been recognized that neurons in the SCN were densely packed (Van den Pol, 1980). However, this 3D reconstruction provides the first demonstration that large soma-soma plate-like contact sites adjoin clusters of core SCN neurons (Fig. 8). The extent of these contact plates distinguishes SCN neurons from those reported in other brain regions, including the cerebral cortex, where extensive studies in mice using both conventional imaging techniques and 3D reconstruction via serial EM have found that neuronal somata typically have no specialized contact sites (Kasthuri et al., 2015; A.N. van den Pol, personal communication, 2019).

**Figure 8.**
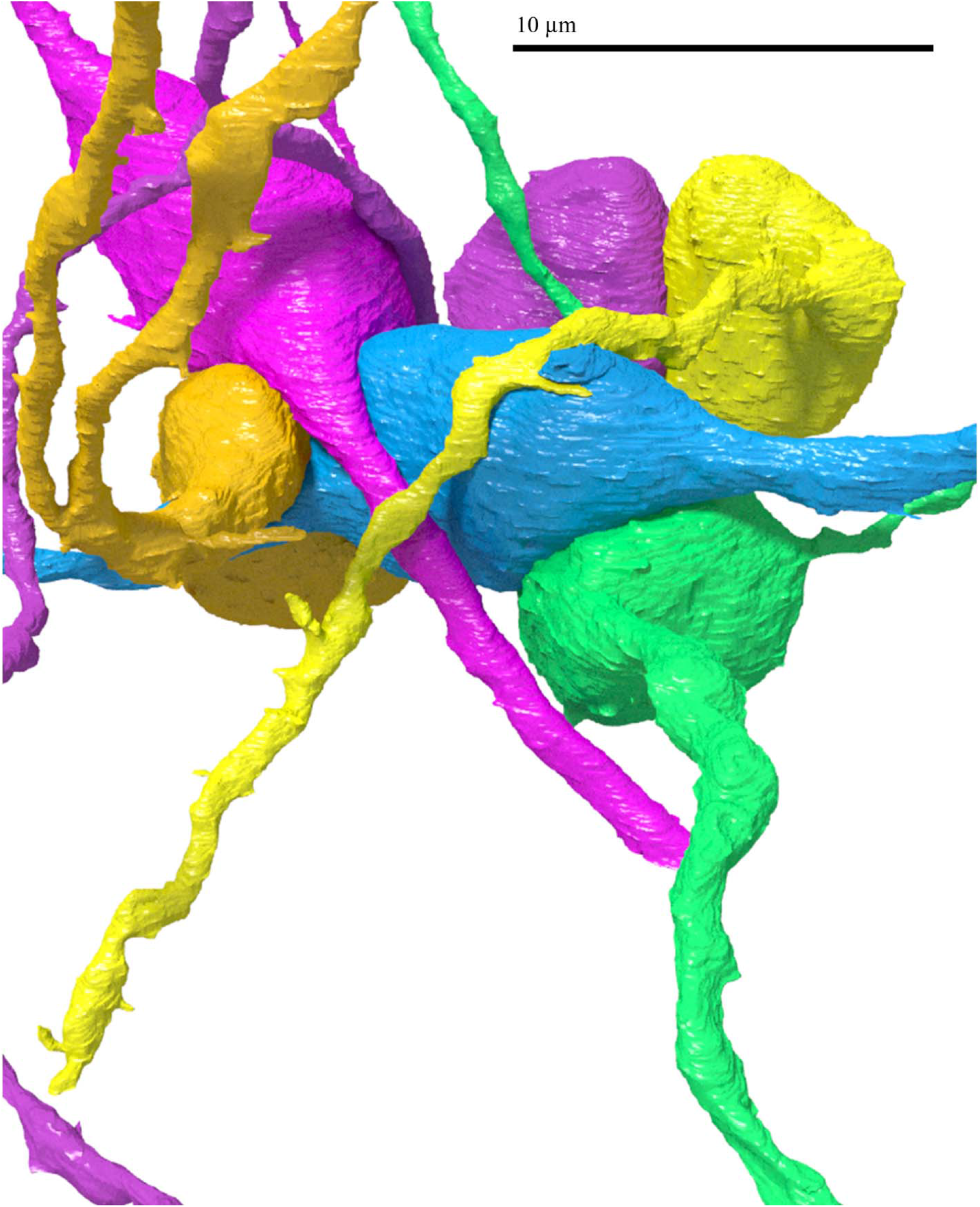
Multiple Soma-Soma Ephaptic Contact Plates Provide Extensive Surface Area Contact Between Clustered SCN Core Neurons. The prominent ephaptic contact plates could subserve circadian synchronization among small-world networks of clustered SCN neurons. The scale bar represents 10 µm.

**Figure 9.**
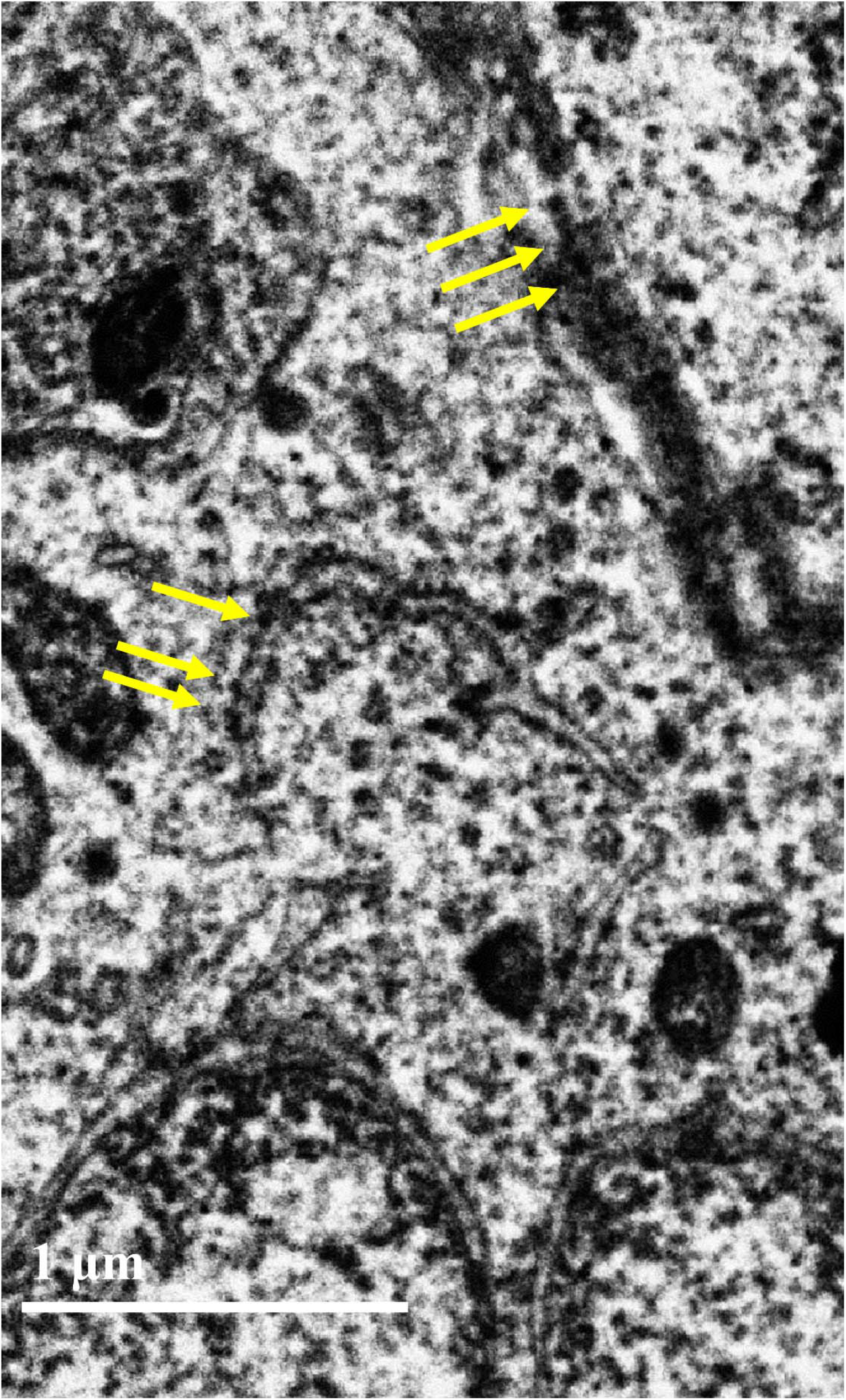
Consecutive, Equally Spaced Punctate Densities Resembling a Zipper Observed Along Adjacent Perikaryal Areas.

The presence of soma-soma plate-like contact sites between intact SCN core neurons is particularly noteworthy, given that dispersed SCN neurons generate desynchronized independent circadian oscillations of spontaneous neuronal firing, whereas neurons in the intact SCN generate synchronized oscillations (Welsh et al., 1995; Herzog et al., 1998). These observations, together with those of Schwartz et al. (1987) demonstrating that *in vivo* synchronization between SCN cells persists for weeks in the absence of action potentials and concomitant synaptic activity, suggest that plate-like contact sites (Fig. 4) may well facilitate synchronization of clustered SCN neurons.

Examination of sequential coronal EM images through six columns within the digitized SCN tissue block revealed that SCN neuronal soma are organized into discrete clusters that are joined not only by dendro-dendritic chemical synaptic networks as previously reported (Kim et al., 2019), but also by plate-like contact sites that are pervasive within this region of the SCN. This finding supports the hypothesis by Yan *et al*. that “SCN network organization may resemble a small-world network” (Watts and Strogatz, 1998; Yan et al., 2010) “…with mostly local connections between nodes….” Dr. Angélique Arvanitaki first proposed that such plate-like contact sites, which she named ephapses [from the Greek *ε*□*απτω*, meaning to touch onto or attach (Arvanitaki, 1942)], are functionally specialized neuronal structures designed to facilitate bidirectional synchronization of spontaneously rhythmic activity in what she termed “pace-maker” neurons (Arvanitaki, 1942; Ogilvie, 1990). Extensive ephaptic plates linking clustered SCN neuronal somata could thus subserve state-independent synchronization of the cell-autonomous oscillators in the SCN that persists even in the absence of synaptic communication, both *in vivo* and *ex vivo*.

Furthermore, electron-dense, dark patches within plate-like contact sites without interleaved glial processes may represent gap junctions embedded within a subset of ephaptic plates. The presence of gap junctions could potentially increase the strength of interneuronal coupling to confer absolute coordination between these self-sustained neuronal oscillators within a given cluster (von Holst, 1939), as has been postulated for cardiac pacemaker cells with similar regions of close intermembrane contact containing electron-dense clusters (perinexal nanodomains) (Hoagland et al., 2019). Conversely, the presence of glial cell membranes nearly devoid of cytoplasm within soma-soma plate-like contact sites form structural barriers to the formation of gap junctions, limiting the potential strength of interneuronal coupling within those regions of the clusters to that of electrical field potentials (Faber and Pereda, 2018). Such weaker coupling could account for the small-world networks that comprise the SCN core, in which local clusters of circadian clock neurons integrate individual rhythms to produce compromise periods among oscillators with disparate periods (Welsh et al., 1995). The results are consistent with von Holst’s description of viscous coupling from more than 80 years ago (von Holst, 1939).

Such synchronization of cortical neurons would likely be incompatible with the functioning of most other brain regions during wakefulness, which may account for the scarcity of observed soma-soma plate-like contact sites in other cortical brain areas. However, during sleep, state-dependent synchronization of cortical neurons also involves even weaker ephaptic coupling between less densely organized neuronal somata in the hippocampus (Chiang et al., 2019), and greatly limits perceptual and sensory processing and other cognitive functions, including consciousness. In contrast, in the SCN, provision of reliable temporal information requires state-independent synchronized neuronal activity, supporting a potentially purposeful function for the strengthened coupling that can be achieved *en masse* by pervasive soma-soma ephaptic contact plates. Ephaptic coupling between SCN neurons in core neuronal clusters—when connected as small-world networks—would enhance the synchronizability of the SCN (Watts and Strogatz, 1998), and thereby improve the functioning of this system as the primary pacemaker for the mammalian circadian clock.

Electric field effects influencing the likelihood of an action potential by affecting membrane potential within a clustered group of neurons could support the maintenance of endogenous rhythmic output and enhance synchronization to the light-dark cycle guided by light-driven input from the retinohypothalamic tract. Photic input from ipRGCs in the optic chiasm to a subset of SCN neurons could be enhanced by ephaptic plate contacts to entrain larger populations of clustered SCN neurons. This may strengthen photic synchronization by co-opting the synchronizing effects of ephapses, and rapidly equip clusters of neurons to relay temporal information. Functional experiments using a different set of techniques would be required to follow up on questions introduced by this work.

The localization of neuronal somata in the optic chiasm, and evidence that processes from some of these neurons extend into the SCN with synaptic connectivity, are noteworthy, because the optic chiasm is largely composed of myelinated axons projecting from the retina to the visual cortex. In addition to synapses within the optic chiasm, retinal axons also enter the ventral part of the SCN, allowing intrinsically photosensitive retinal ganglion cells to innervate SCN neurons, potentially providing the non-image-forming light-dark information required to entrain the SCN pacemaker to external light-dark information (Paul et al., 2009). Very few neurons reside in midline structures in the SCN (Jeffery, 2001), yet the reconstructed pair of neurons with somata in the optic chiasm adjacent to the SCN core are two of several dozen neurons interspersed among the optic nerve fibers crossing over the midline. These reconstructed cells could thus have received photic input from ipRGCs that served as external synchronizers from environmental light-dark cycles. This would be consistent with an immunocytochemical analysis of neurons in the rat optic chiasm reporting VIP-containing neuronal somata found entirely embedded in the optic chiasm adjacent to the SCN core (Card et al., 1981). A second possible explanation for the presence of these neurons in the optic chiasm is that they may be components of an interhemispheric bridge between the two suprachiasmatic nuclei (1999).

The findings of this report are subject to limitations. First, this SCN reconstruction is from a small region within the SCN of a single young adult nocturnal male mouse prepared at one time of day (ZT 10). The SCN is sexually dimorphic (Bailey and Silver, 2014), changes structurally at different developmental stages (Kabrita and Davis, 2008), and undergoes fundamental network reorganization across circadian phases (Yan et al., 2010). Further SEM 3D reconstructions of the entire SCN at different developmental stages and across circadian time in sexually diverse mammals would therefore be required to characterize the SCN connectome comprehensively.

Second, although neurons were intentionally selected from deep in the block to maintain maximal dendritic extent, dendrites were truncated at the boundaries of the region of interest, excluding distal dendrites from the structural analysis. However, it is unlikely that processes far from the cell body would contribute as strongly to local synchronization as proximal structures. Third, recent studies have discovered distinctive roles of different SCN cell types in the structural (Wen et al., 2020) and functional (Shan et al., 2020) connectome. Although of interest, specific cell types were not identifiable in this 3D-reconstruction. Thus, while we examined the SCN core, where about 25% of the neurons contain VIP, we cannot determine whether neurons of different chemical types in the SCN have different patterns of connectivity. Future studied employing genetic labeling of VIP or other cell types would help to elucidate ultrastructural characteristics of specific cell types. Nonetheless, we found tight clustering of SCN neurons to be pervasive within the image volume. Fourth, though the sections were ultrathin (50-nm), the thickness of each section limited the resolution possible in the Z-dimension relative to the 4 × 4-nm-per-pixel resolution in the X- and Y-dimensions. Future reconstructions of the SCN could focus on sectioning in a different orientation (e.g., sagittal or horizontal) to overcome this limitation. Advancing technologies in the field of connectomics are being developed to expedite large-scale SEM data collection and analysis with possibilities including genetically labeled cell types and correlative light electron microscopy (Joesch et al., 2016; Fang et al., 2018). Parallel studies could explore evidence for ephaptic coupling by characterizing the phase relationships among tightly clustered SCN neurons and generating distance-based phase-prediction models.

In summary, using high-resolution images of 805 serial ultrathin sections from a scanning electron microscope, we created digitized three-dimensional reconstructions of mammalian SCN neurons, which revealed distinct ultrastructural morphology of SCN neurons. The morphology included tight clustering of SCN neurons and soma-to-soma plate-like contact sites with and without interleaved glial processes, which may mediate ephaptic coupling necessary for the tissue-wide synchronization of pacemaker neurons with disparate intrinsic periods within the SCN core. These features could potentially provide a non-synaptic mechanism—via ephaptic interactions—for the synchronizing communication between proximate core SCN neurons required to generate the synchronous oscillations. Moreover, these results demonstrate an effective and reproducible pipeline for cutting and imaging ultrathin serial sections of the mouse SCN for 3D reconstruction at ultrastructural resolution, which can guide future high-resolution imaging experiments in the primary organism used for modeling circadian clock synchronization.

## Conflicts of Interests

The authors declare no competing financial interests.

## Acknowledgements

Research was supported by grants from the National Institute of Mental Health (NIMH; P50 MH094271) and National Institute of Neurological Disorders and Stroke (NINDS; U24 NS109102). We thank Dr. Xueying (Snow) Wang at Harvard and Mrs. Chrysanthi Joseph at UT Southwestern for assistance with the mouse transcardial perfusion. We further thank Dr. Clifford Saper for reviewing the manuscript and providing informative feedback.

## Author Contributions

Study concept: MÉC, JWL, JST. Study design: MÉC, JWL, JST. Acquisition, analysis and interpretation of data: MÉC and DRB. Drafting of the manuscript: MÉC. Critical revision of the manuscript for important intellectual content: YS and DRB. Technical and material support: YS, RS, DRB, AS-P, JWL, JST. Scientific direction and funding: JST, JWL.

## Multimedia Legends

**Multimedia Video 1**. Layers of Reconstruction via Volume Annotation and Segmentation Tool. Seconds 1-4 show 125 consecutive serial sections containing EM image data. Seconds 5-8 show 125 consecutive serial sections containing EM images with the segmentation layer containing manually traced and color-labeled objects. Parts of the neurons assigned blue, yellow, green, orange, pink, and purlple are shown. Seconds 9-13 show 150 consecutive serial sections containing the segmentation layer without the background EM images. Frames were collected as screenshots from VAST and rendered at 30 frames per second. <u>View Multimedia Video 1 online</u>.

**Multimedia Video 2**. Rotation and *in silico* Dissection of Cluster of Six Reconstructed Somata Highlights Multiple Large Reciprocal Soma-Soma Ephaptic Contact Plates Linking SCN Core Neurons. The neurons shown in the video retain their assigned color from the initial reconstruction (blue, yellow, green, orange, pink, and purple). Each point of ephaptic contact between each neuron and an adjacent soma was labeled in a different, previously unused color (magenta, dark green, tan, lime green, pale green, and dark red). The video shows a looping rotation of the somata as positioned in the SCN core, and then pulled apart to reveal ephapses between these somata. Soma-dendrite ephapses are not shown. Video was animated in 3ds Max and rendered at 30 frames per second. <u>View Multimedia Video 2 online</u>.

**Multimedia Video 3**. Overview Video Shows Perspectives of the Eight Reconstructed SCN Neurons Superimposed on the First Serial Section. Using a camera to approach, zoom in on, and rotate around the eight neurons highlights architecture that could support ephaptic (soma-soma, soma-dendrite ephapses) and synaptic transmission (axons and dendrites extending in the direction of different regions of the SCN and optic chiasm). Together, ephaptic and synaptic coupling could support all three levels of synchronization (internal synchronization of the small-world, clustered networks, receiving input from the optic chiasm, and extending outside of clusters via synaptic and ephaptic connections). The EM image of the serial section provides a way to distinguish between the SCN and optic chiasm regions, as evidenced by the region composed primarily of cells (the SCN, which contains the somata of the color-labeled blue, yellow, green, orange, pink, and purple neurons) versus the region composed primary of myelinated axons (the optic chiasm, which contains the somata of the color-labeled navy and red neurons). Video was animated in 3ds Max and rendered at 30 frames per second. <u>View Multimedia Video 3 online</u>.

**Multimedia Videos 4-10**. Columnar Spacelapse Videos Display the Pervasiveness of SCN Neuronal Soma-Soma Ephapses. Each video shows a randomly selected spacelapse recording from the first section through the 805^th^ section capturing a different window-of-view from the EM-imaged ROI. In each video, there are clustered neuronal somata contacting each other through ephaptic plates, demonstrating that the six neurons reconstructed from the SCN with ephaptic plates are representative of clustered neurons throughout the SCN tissue collected.

Videos were produced by compiling screenshots of each consecutive section within the window-of-view in VAST and rendered at 35 frames per second. Refer to Figure 8 to identify the window-of-view from which each video was generated.

<u>View Multimedia Video 4 online</u> under the channel Mark Czeisler.

<u>View Multimedia Video 5 online</u> under the channel Mark Czeisler.

<u>View Multimedia Video 6 online</u> under the channel Mark Czeisler.

<u>View Multimedia Video 7 online</u> under the channel Mark Czeisler.

<u>View Multimedia Video 8 online</u> under the channel Mark Czeisler.

View Multimedia Video 9 online under the channel Mark Czeisler (a link will be made available at a later time).

<u>View Multimedia Video 10 online</u> under the channel Mark Czeisler.

## Notes

### Competing Interest Statement

The authors have declared no competing interest.

